# SHARK: a specialized host for assembling R6K plasmids

**DOI:** 10.1101/2025.10.06.680659

**Authors:** Shivang Hina-Nilesh Joshi, Christopher Jenkins, David Ulaeto, Thomas E. Gorochowski

**Affiliations:** School of Biological Sciences, University of Bristol, 24 Tyndall Avenue, Bristol, BS8 1TQ, UK; CBR Division, Defence Science and Technology Laboratory, Porton Down, Wiltshire, SP4 0JQ, UK

**Keywords:** Pir strains, R6K plasmids, conditional replication, cloning, lambda-RED

## Abstract

R6K plasmids are commonly used for a wide range of genome engineering applications due to their ability to support transient delivery of genetic cargoes in many hosts. The maintenance of R6K plasmids requires specific strains. Unfortunately, many of these have obscure backgrounds, limited availability and were not built for efficient cloning. To address this issue, we present the construction and characterisation of a series of Pir *E. coli* strains called SHARK that are built from the DH10B derivative, Marionette-Clo. All SHARK strains have a genome encoded Pir gene for stable R6K plasmid maintenance and a *λ*CI gene for tight unconditional repression of specific genes on plasmids. We show that SHARK strains are *>*100-fold more efficient than a commercial Pir strain for large and complex cloning reactions. SHARK is intended to help facilitate the cloning of large and complex R6K plasmids for challenging genome engineering projects, with all strains and genetic tools for their assembly being made publicly available.

**TABLE OF CONTENTS IMAGE:** 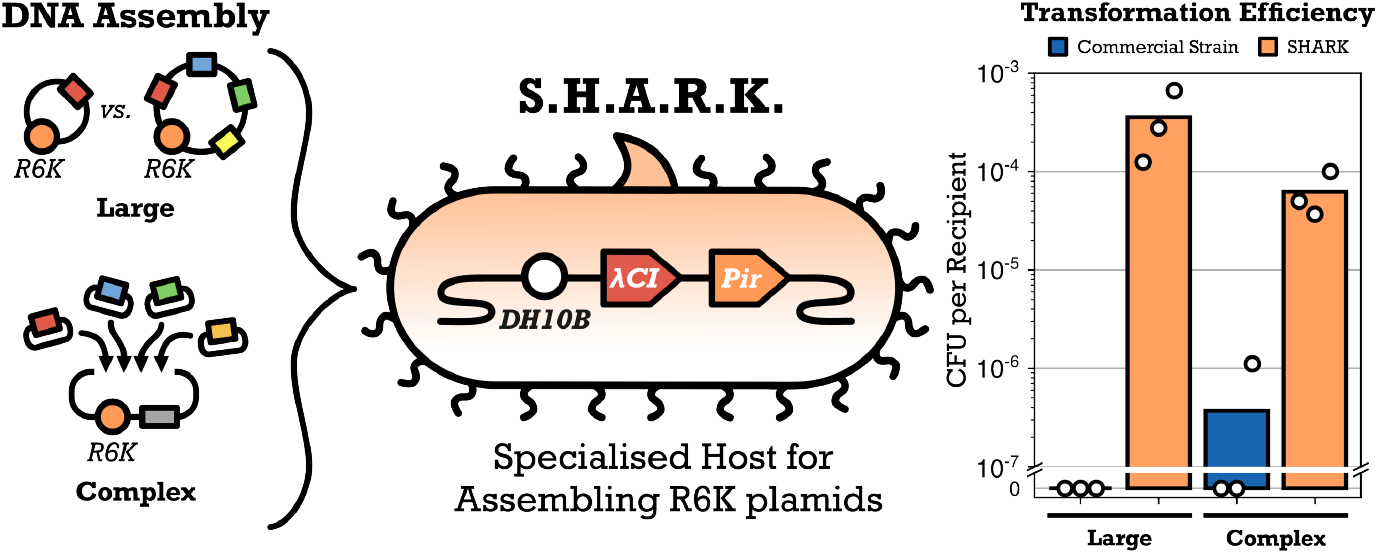

## INTRODUCTION

R6K vectors are widely used for microbial genome engineering and have been employed for homologous recombineering [1, 2], phage recombinase integration [3–5], and random transposon insertion [6–8]. They have also been instrumental in delivering a variety of DNA cargoes including genome landing pads [9, 10], small molecule sensor arrays [11], genetic logic circuits [12], recombinase-based memory circuits [13], and metabolic pathways [5]. This widespread use stems from their inability to replicate outside of specific cloning strains, enabling precisely controlled transient delivery of a DNA cargo.

All R6K vectors are derived from the *γ*-origin of the wildtype R6K plasmid [14] where the vegetative region, oriV (A+T rich region with 7 direct repeat sequences), is present on the plasmid and the Pi replication gene, *pir*, is integrated into the host genome (**Figure 1a**). By providing the Pir protein in trans, plasmid replication is able to initiate at oriV and ensure plasmid maintenance. This wildtype system is autoregulatory where the Pir protein forms homodimers at high concentrations, which leads to it no longer being able to activate plasmid replication. Therefore, expression of the Pir protein has to be carefully regulated within these strains. To overcome this issue, “copy-up” mutants of the Pir protein have been created that do not display this inhibition and can maintain R6K vectors at elevated copy numbers [15, 16].

**Figure 1:**
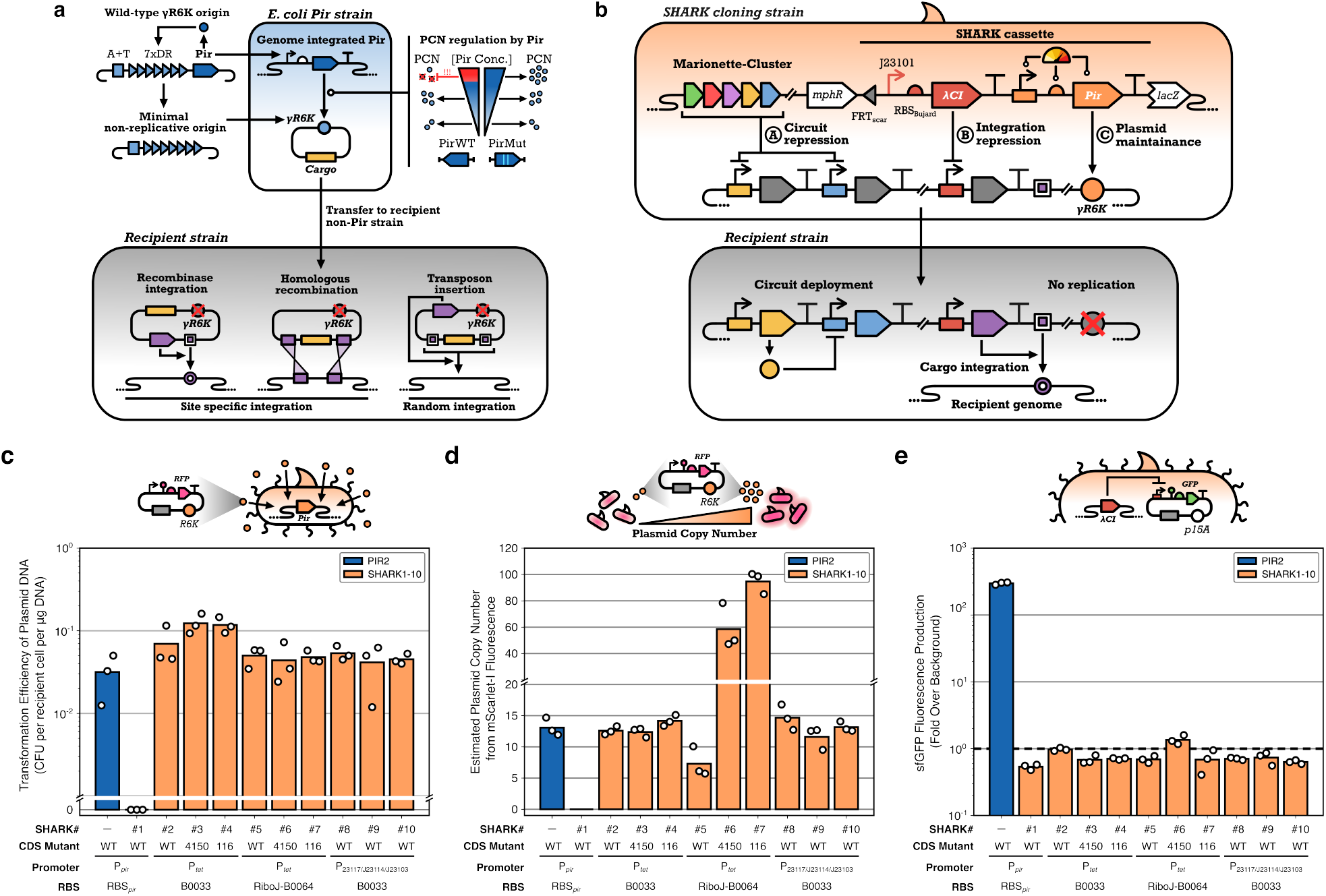
Overview of the SHARK strain design. (**a**) Non-replicative R6K vectors are derived from the *γ* origin of replication of the wildtype R6K plasmid. This origin consists of a A+T rich region (DNA replication initiation site), 7 direct repeat (7xDR, Pir monomer binding sites), and a auto-regulating Pir gene. The vegetative components of *γ*R6K forms the minimal non-replicative origin, which can only be maintained in *E. coli* strains with a genomically integrated Pir gene. Mutant Pir genes have also been used that elevate plasmid copy number. Genetic circuit schematics shown using Synthetic Biology Open Language (SBOL) Visual symbols [41]. (**b**) SHARK strains are built on the Marionette-Clo strain, a DH10B derivative with 12 evolved inducible transcription factors in the genome. SHARK strains express a Pir protein to maintain R6K plasmids, and a *λ*CI protein to repress plasmid genes (e.g., those involved in plasmid integration into the genome). (**c**) Transformation efficiency PIR2 and SHARK strains with purified miniprepped plasmid DNA with a *γ*R6K origin. (**d**) Estimated R6K plasmid copy number in PIR2 and SHARK strains. (**e**) sfGFP fluorescence from a plasmid where sfGFP protein expression is driven by a *λ*-promoter from a p15A plasmid. Dashed line denotes cell autofluorescence level. PIR2 results shown in blue, SHARK results shown in light orange.

Numerous specialized *E. coli* strains, simply called Pir strains, have been created for R6K plasmid maintenance. These can be purchased commercially (e.g., Invitrogen One Shot PIR series and Lucigen Transfor-Max EC100D series), are available from culture collections (e.g. BW29427, S17–1*λ*pir[17], MFDpir[18], *HB101–λ*pir[19]), or can be directly requested from research groups (e.g., DH5Apir [6], CC118/MV1190 [20], DIAL[21]). However, many of them were developed several decades ago from base strains with unclear backgrounds and their performance for typical cloning tasks is not known. For example, we have personally observed low transformation efficiencies and plasmid instability when using a commercial Pir strain (PIR2) for cloning genome engineering plasmids [4]. In contrast, no such issues were seen for another *E. coli* strain, DH10B, which we use commonly for complex cloning tasks [22, 23].

In this work, we address this issue by developing a set of DH10B derived Pir strains called SHARK (Specialized Hosts for Assembling R6K plasmids). SHARK strains are specifically built from Marionette-Clo [11], an *E. coli* DH10B derivative with 12 inducible transcription factors integrated into its genome. To adapt this strain for working with R6K plasmids, we genome integrated a variety of Pir expression cassettes that explored different combinations of promoter and ribosome binding site (RBS) strength, as well as the Pir protein coding sequence. We find that the vast majority of our strains are able to successfully maintain R6K plasmids, and comparisons of one of our best performing strains (SHARK2) to a commercial variant (PIR2, Invitrogen) showed that transformation efficiencies of plasmid assembly reactions in SHARK2 were *≥*132-fold higher. Furthermore, our SHARK strains contain a constitutively expressed *λ*CI gene, thereby providing unconditional and tight repression of genes commonly found on genome engineering plasmids, reducing the chance of activation of these systems during cloning. Overall, our results show that SHARK strains can support the cloning of larger and more complex R6K plasmids for diverse applications requiring the transient delivery of genetic cargoes.

## RESULTS

### Design and construction of SHARK strains

As a foundation for our SHARK strains, we opted for the widely used *E. coli* DH10B strain due to its demonstrated track record as an efficient cloning strain with a high transformation efficiency and ability to work with large plasmids [22, 23]. In particular, we built all of our SHARK strains from Marionette-Clo [11], a derivative of DH10B that also contains a set of 12 inducible transcription factors integrated into its genome. This allows for repression of a broad range of cargoes and helps to reduce the burden imposed on the host when cloning large plasmids containing complex genetic circuitry [24, 25] **(Figure 1b)**.

To ensure stable maintenance of R6K plasmids in our strains, it is essential that the Pir protein is sufficiently expressed. However, Pir cannot simply be over-expressed as this could result in protein dimerization and a breakdown in plasmid replication [26, 27]. Optimization of Pir expression was therefore critical for robust R6K plasmid maintenance. As other Pir strains have been made using the wildtype promoter and RBS sequence [15], we employed this design for SHARK1 by retrieving these sequences from a previous study [28]. In addition, we built a number of synthetic designs, SHARK2–7, where we varied features of the Pir transcription unit to systematically vary its expression and reduce our reliance on uncharacterised native regulatory components. We specifically used an inducible P_*tet*_ promoter[29] to enable controlled regulation of transcription (the cognate TetR repressor is provided from the Marionette-Clo genome) and tested a weak B0033 and strong B0064 RBS to regulate translation, with the B0064 RBS also including a RiboJ self-cleaving ribozyme that has been shown to boost translation levels through both stabilization of transcripts and a reduction of structure around the RBS [30]. We also considered copy-up mutants of the Pir protein coding sequence, including the moderate mutant Pir4150 (T108I and P113S) [16] and the more frequently used Pir116 (P106L) [15] to see if copy number of R6K plasmids could be controlled.

Each of these different designs for expressing the Pir protein were encoded into SHARK cassettes (**Figure 1c**) and placed in pBBR1 plasmids with a FRT-flanked spectinomycin resistance marker to allow for selection and easy removal of the marker using Flp recombinase after genome integration (**Supplementary Figure 1**). In addition, each cassette also included a *λ*CI coding sequence expressed from a strong constitutive promoter (P_J23101_) and strong ribosome binding sequence (RBS_Bujard_) to provide robust repression of plasmids containing genome integration machinery whose expression is driven by a *λ*-promoter [4].

Once these SHARK cassette plasmids had been built, their cargoes were integrated into the genome of Marrionette-Clo using the *λ*-RED system [31]. This system is known to be efficient for small payloads and can be customized to target any site for insertion using short homology arms that are easily added using PCR (**Supplementary Figure 1**). To simplify this process, we created our own *λ*-RED plasmid-based system called pREMORAv1 that was used to build every SHARK strain in this study. The pREMORAv1 system consists of a salicylic acid inducible *λ*-RED operon encoding the *β, γ* and exo genes, and a constitutively expressed *recA* gene, on a low-copy plasmid with a temperature sensitive pSC101 origin of replication and beta-lactamase selection marker. We targeted the insertion of the SHARK cassettes to the wildtype *lacI* locus, and post integration, all strains were cured of the spectinomycin selection marker using transient expression of Flp recombinase so that the final SHARK strains were marker-free.

To validate that the SHARK strains were able to maintain R6K plasmids, all designs were made chemically competent and transformed with pR6K-mScarI, a *γ*R6K plasmid constitutively expressing the mScarletI red fluorescent protein (**Supplementary Figure 3**). We were initially surprised to find that SHARK1 containing the wildtype promoter, RBS and Pir gene could not be transformed with pR6K-mScarI. However, upon closer examination of the full sequence of the Pir gene from the original study [28], we found that the oligo used in later work to create the uidA-pir fusion [15] was missing 4 nucleotides from the 5^*′*^ of the upper Pir binding inverted repeat (IR) sequence. This may weaken the auto-regulation of the Pir gene, resulting in low levels of Pir protein expression. In SHARK1, we used the complete P3 Pir promoter sequence from the original study [28], which has both intact IR sequences, and this may possibly impose tighter repression of Pir expression.

In our other designs, SHARK2–7, synthetic parts were used to regulate Pir expression and we further tested different copy-up mutants of the Pir protein coding sequence. We found that all these designs could be transformed with pR6K-mScarI. Interestingly, we also discovered that none required anhydrotetracycline hydrochloride (aTc) induction of Pir expression for plasmid maintenance, suggesting that low basal levels of expression from the P_*tet*_ promoter was sufficient for R6K plasmid maintenance.

As the inducible function of the P_*tet*_ promoter was found to serve no purpose, we decided to create a set of additional strains, SHARK8–10, where P_*tet*_ was swapped for one of three weak constitutive promoters (P_J23117_, P_J2314_ and P_J23103_). This would allow the TetR system to then be freed up for other purposes. We found that all of these new designs could be successfully transformed and were able to maintain the pR6K-mScarI plasmid (**Figure 1c**).

### Estimating R6K plasmid copy number in SHARK strains

To assess the stability of R6K plasmid replication within our strains, we used fluorescence to estimate plasmid copy number [32, 33] (**Supplementary Figures 3 and 4**). Specifically, we compared the relative fluorescence of a mScarlet-I gene from an R6K plasmid (pR6K-mSarI) against a p15A plasmid (p15A-mScarI), which has a well known copy number of 9 copies per genome [32–34] (**Methods**). To improve accuracy, the mScarlet-I gene was designed to be weakly expressed so that the measured signal would be limited by DNA copy number.

Using this method, we estimated that the commercial PIR2 strain maintains pR6K-mScarI at approximately 13.1 copies per genome. This closely matches the expected 15 copies per genome reported in the literature [15]. Even with no addition of aTc, most of the SHARK strains using P_*tet*_ to express the Pir protein were able to maintain R6K plasmids at low to medium copy numbers, with SHARK2–4 and SHARK8–10 having estimated copy numbers in the range of 11.6–14.7 copies per genome (**Table 1**), similar to the PIR2 strain. In contrast, SHARK5 maintained R6K plasmids at a slightly lower copy number of 7.3 copies per genome, and SHARK6 and SHARK7 maintained plasmids at elevated copy numbers of 58.5 and 94.7 copies per genome, respectively. In all cases, we found that R6K plasmids were replicating at a sufficient rate for stable propagation.

**Table 1:** Strain designs and their estimated R6K plasmid copy numbers^a^.

| Strain | Pir expression cassette |  |  | R6K PCN<sup>b</sup> |
| --- | --- | --- | --- | --- |
|  | Promoter | RBS | Protein |  |
| PIR2 | P<sub>pir</sub> | RBS<sub>pir</sub> | Pir | 13.1 ± 1.9 |
| SHARK1 | P<sub>pir</sub> | RBS<sub>pir</sub> | Pir | — |
| SHARK2 | P<sub>tet</sub> | B0033 | Pir | 12.6 ± 0.7 |
| SHARK3 | P<sub>tet</sub> | B0033 | Pir4150 | 12.4 ± 0.8 |
| SHARK4 | P<sub>tet</sub> | B0033 | Pir116 | 14.2 ± 0.9 |
| SHARK5 | P<sub>tet</sub> | RiboJ-B0064 | Pir | 7.3 ± 2.4 |
| SHARK6 | P<sub>tet</sub> | RiboJ-B0064 | Pir4150 | 58.5 ± 17.2 |
| SHARK7 | P<sub>tet</sub> | RiboJ-B0064 | Pir116 | 94.7 ± 8.3 |
| SHARK8 | P<sub>J23117</sub> | B0033 | Pir | 14.7 ± 2.0 |
| SHARK9 | P<sub>J23114</sub> | B0033 | Pir | 11.6 ± 1.8 |
| SHARK10 | P<sub>J23103</sub> | B0033 | Pir | 13.2 ± 0.8 |

a. PIR2 is a commercially available Pir strain from Invitrogen. The P<sub>pir</sub>, RBS<sub>pir</sub> and Pir regulatory parts and coding sequence are wildtype sequences from the original study [15]. Full part sequences are provided in **Supplementary Data 1**.

For SHARK strains where the Pir protein was expressed from a P_*tet*_ promoter, we also assessed how R6K plasmid copy number might be affected after P_*tet*_ induction with aTc (**Supplementary Figure 4**). For SHARK2–4, induction with aTc did not affect plasmid copy number, possibly because the B0033 RBS is very weak. In contrast, mixed results were seen for SHARK5–7, which contained a stronger RBS. For SHARK5, which contains the wildtype Pir protein coding sequence, induction with aTc resulted in the loss of the plasmid from the cells. This is expected as others studies that have shown excessive expression of the wildtype Pir protein causes inhibition of plasmid replication [14]. For SHARK6 and SHARK7, which contain copy-up mutants of the Pir protein, induction with aTc caused a notable increase in mScarlet-I fluorescence of approximately 1.6-fold for SHARK6 and 1.75-fold for SHARK7 when comparing uninduced and fully induced (100 nM aTc) samples. Large variability in plasmid copy number was seen across biological replicates for SHARK6, while much more reproducible and statistically significant increases in plasmid copy number were seen for SHARK7. Overall, SHARK7 was the only strain that saw reliable control of R6K plasmid copy number, allowing it to be varied from approximately 95 to 166 copies per genome.

### Genome integrated *λ*CI efficiently represses gene expression from plasmids

A common use case for R6K plasmids is the transient expression of genetic cargoes for genome integration. Many of the plasmids designed for this task make use of temperature sensitive *λ*CI proteins to enable controlled expression of specific components, e.g., integrase enzymes that integrate the entire plasmid at a target locus. While working with such systems, like the widely used one-step integration plasmid (OSIP) [4], we have seen undesired integration of cargoes into cloning strains even though a temperature sensitive *λ*CI gene is present on the plasmid and cells are cultured at 30°C. This can be problematic as it makes the plasmid selection marker redundant and destabilizes the plasmid.

To overcome this issue, our SHARK strains include a constitutively expressed *λ*CI gene to ensure strong repression of *λ*-promoters. To confirm repression by the genome integrated *λ*CI in our strains, all SHARK strains were transformed with a p15A plasmid containing a *λ*CI regulated promoter driving sfGFP expression and green fluorescence was measured from cultures. We found that all SHARK strains could successfully repress sfGFP expression to near cell autofluorescence levels (**Supplementary Figure 5**). Of note, despite the Pir gene being non-functional in SHARK1, the *λ*CI gene functions just as well as in the other SHARK strains. To demonstrate this further, we transformed SHARK1 with a genome integrative plasmid and observed no genome integration, even in conditions where cells were incubated overnight at 37°C, where the plasmid encoded *λ*CI_*ts*_ does not contribute to recombinase inhibition (**Supplementary Figure 6**). This demonstrates that the genome encoded *λ*CI is sufficiently expressed to fully inhibit even transient expression from the plasmid.

### SHARK is an efficient cloning strain for large and complex DNA assemblies

The assembly and cloning of plasmids is often a bottleneck in synthetic biology projects. The difficulty of assembling a construct can vary depending many factors, such as the method of assembly, plasmid size, number of parts, as well as by the contents of the assembly itself. To benchmark the performance of our SHARK strains for common cloning tasks, we transformed SHARK2 and the commercially available PIR2 strain with a set of four different assembly reactions: a (i) 4 kb and (ii) 8 kb blunt-end ligation (BEL) reaction, and a (iii) 2-part and (iv) 5-part Golden-Assembly (GGA) reaction. These assemblies are representative of the features that typically affect assembly efficiency, i.e., complexity (2-part vs. 5-part), plasmid size (4 kb vs. 8 kb), and contents of assembly (no cargo vs. inducible phage RNAP insertion). All of the assemblies tested used the pOSIP-CH plasmid [4] as a backbone. This is a genome integration vector containing a R6K origin or replication and a recombinase gene whose expression is regulated by a *λ*CI repressor.

Experiments showed that SHARK2 had a higher transformation efficiency than PIR2 for all assembly reactions tested. For the simpler assembly reactions (4 kb BEL and 2-part GGA), SHARK2 had approximately 484-fold and 132-fold higher transformation efficiencies than PIR2. For the more difficult assemblies (8 kb BEL and 5-part GGA), PIR2 did not produce any transformants for most of the transformation reactions, while numerous transformants were seen for SHARK2. Comparing the simple and difficult assemblies for SHARK2 alone, there was a notable 6.7-fold drop in transformation efficiency between 4 kb vs 8 kb BEL assemblies, and a 20-fold drop between 2-part GGA and 5-part GGA, highlighting the increased challenge of working with these larger and more complex assemblies.

## DISCUSSION

In this work, we have presented the design and construction of a series of *E. coli* Pir strains called SHARK that are capable of maintaining and being efficiently transformed with R6K plasmids. Out of the 10 SHARK strains constructed, 9 were capable of robust maintenance of R6K plasmids at either low/medium (SHARK2–5 and SHARK8–10) or high (SHARK6–7) copy numbers (**Figure 1**), and we showed that one of our best performing strains (SHARK2) could be effectively transformed with large and complex plasmid assembly reactions that often lead to no colonies in a commercially available Pir strain (**Figure 2**). All SHARK strains also included a genome integrated *λ*CI gene, which we showed was capable of effectively silencing plasmid-based gene expression from *λ*-promoters.

**Figure 2:**
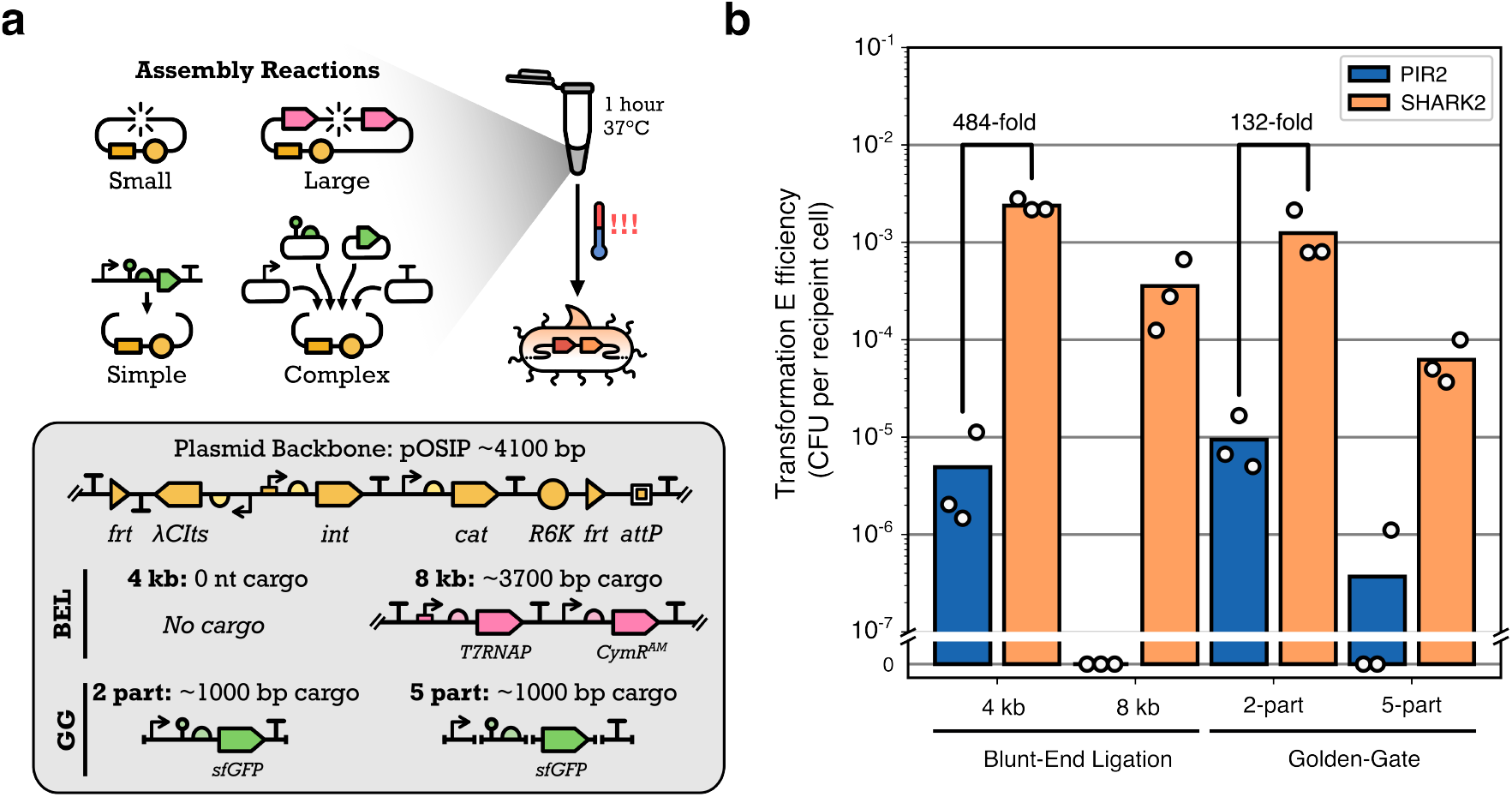
Transforming SHARK2 and the commercial PIR2 strain with cloning assembly reactions. (**a**) SHARK2 and PIR2 were transformed with four cloning assembly reactions testing a combination of assembly sizes (4 kb and 8 kb) and assembly complexity (2-part and 5-part) for two different types of cloning reactions (Blunt-End Ligation, BEL, and Golden-gate assembly, GGA). All the assembly reactions were performed using the pOSIP-CH plasmid backbone, a suicide vector designed for *E. coli* genome engineering. (**b**) Transformation efficiency of SHARK2 and the commercial PIR2 strains were calculated and reported as CFU per recipient cell. PIR2 results shown in blue, SHARK2 results shown in light orange.

Given our results, we recommend SHARK2 or SHARK8 for routine cloning tasks. Both these strains have identical designs, except the promoters driving expression of the Pir protein are P_*tet*_ and P_J23117_, respectively. SHARK2 and SHARK8 maintain R6K plasmids at medium copy numbers of 12.6 and 14.7 copies per genome, respectively, which is useful when cloning large constructs, or if the plasmid expresses burdensome or toxic genes. If higher plasmid yields are required, we recommend SHARK6 or SHARK7, as these strains maintain plasmids at higher copy numbers of 58.5 and 94.7 copies per genome, respectively, with copy number also able to be increased by 1.75-fold through further induction with aTc (**Supplementary Figure 3**). A by product of this work was the finding that very little Pir protein is required for robust replication of R6K plasmids. Even basal expression from the P_*tet*_ promoter and the use of a weak B0033 RBS was sufficient to produce a sufficient concentration of Pir protein within a cell (**Figure 1d**).

A key benefit of using SHARK over other Pir strains is its ability to handle complex and large plasmid assemblies. We demonstrated this using a variety of plasmid assembly reactions and showed over 2-orders of magnitude higher efficiencies for SHARK2 over the commercially available PIR2 strain (**Figure 2**). This efficiency is essential when working with large combinatorial libraries, as well as large multi-unit genetic circuits. Furthermore, these types of application are seeing growing interest, with R6K vectors being used for creating cross-species genetic libraries [35, 36], and even cross-kingdom applications [37, 38]. Beyond the use of R6K plasmids as a host for transient genetic cargoes, the small size of the *γ*R6K origin (~390 bp) and its lack of any function once delivered to a target cell, makes these plasmids an ideal foundation for testing synthetic origins of replication. For example, using LLMs to generate potential sequences [39], which can then be tested to create new tools or uncover mechanisms of DNA replication and feedback control.

As interest continues to grow around genome engineering and the integration of large and complex genetic circuits into a wider variety of organisms, our SHARK strains provide a robust and versatile chassis to support these ambitions, overcoming existing plasmid assembly and propagation bottlenecks.

## METHODS

### Strains and media

Initially, *E. coli* NEB 10-beta cells (New England Biolabs, C3019H) were used for cloning all non-R6K plasmids, i.e., SHARK cassette plasmids (**Supplementary Figure 1a**), and One Shot PIR2 cells (Invitrogen, C111110) were used for cloning R6K plasmids, i.e., pR6K-mScarI used in (**Figure 1d–e**). Once SHARK2 was created and its function confirmed, it was then used for all subsequent cloning in this work. All SHARK strains are a derivative of Marionette-Clo (strain sAJM.1504), which was a gift from Christopher Voigt (Addgene plasmid No. 108251) [11]. Lysogeny Broth Miller (LB; Sigma-Aldrich L3522) was used for routine culture preparation for plasmid DNA extractions using the Monarch Spin Plasmid Miniprep Kit (New England Biolabs, T1110); LB Miller with agar (Sigma-Aldrich, L3147) was used for all solid media cultures; Super Optimal Broth (SOB; VWR J906-500G) was used for cell cultures when preparing heat-shock chemical competent cells; M9-glucose (1X M9 salts, Sigma-Aldrich M6030; 0.25 mg/mL thiamine hydrochloride, Sigma-Aldrich T4625; 2 mg/mL casamino acid, VWR J851; 4 mg/mL glucose, Sigma-Aldrich G7528; 2 mM magnesium sulfate, Sigma-Aldrich M2643; 100 µM calcium chloride, Sigma-Aldrich C1016) was used for all fluorescence quantification experiments. Antibiotics were used at the following concentrations: Kanamycin (Kan; Sigma-Aldrich, K1637) 50 µg/ml, Chloramphenicol (Cam; Sigma-Aldrich C0378) 34 µg/ml, Spectinomycin (Spec; Sigma-Aldrich S4014) 50 µg/ml and Carbenicillin (Carb; Sigma-Aldrich C1389) 100 µg/ml. All antibiotics were stored as 1000X working stocks in either 50% glycerol (Kan, Spec, Carb) or 100% ethanol (Cam). Anhydrotetracycline hydrochloride (aTc; Merk 37919) was used to induce SHARK2–7 **(Supplementary Figure 3)** at the following final concentrations: 0, 1, 2.5, 5, 10, 25 and 100 nM. Salicylic acid (Salicylate; Sigma-Aldrich, 247588) was used to induce the *λ*RED operon at a final concentration of 100 µM.

### Heat-shock chemical competent cell preparation

All heat-shock chemical competent (HSCC) cells were prepared in SOB media, and all 37°C incubation steps were done in a New Brunswick Innova 42 shaking incubator (Eppendorf, M1335-0002). Competent cell batches were prepared in either 5 mL volumes in 14 mL polystyrene culture tubes (StarLabs, I1485-2810) for **Figure 1d–e**, 50 mL volumes in 250 mL non-baffled conical flasks for **Figure 2b**, or 200 mL cultures split across four 250 mL non-baffled conical flasks with 50 mL cultures each for **Supplementary Figure 1a**. For as high efficiency as possible, it is best to keep tubes and reagents on ice as much as possible throughout the protocol. Specific instructions for *λ*RED competent cell preparation are detailed later.

Regardless of volume size, the HSCC protocol was as follows. Strains to be made competent were streaked out from glycerol stocks onto non-selective LB agar plates and incubated overnight at 37°C. The following day, a single colony for each strain was inoculated into 2 mL of SOB in 14 mL polystyrene culture tubes and incubated overnight at 37°C shaking at 250 rpm. The following day, the overnight cultures were diluted 1:1000 into 5, 50 or 200 mL of fresh SOB media and cultured for 3 hours at 37°C, shaking at 250 rpm. At 3 hours, the cultures were placed in a ice bath for 20 min; the large *≥*50 mL cultures were aliquoted to 50 mL tubes then placed in the ice bath. The cell cultures were then pelleted in a pre-chilled centrifuge (Eppendorf, Centrifuge 5910 Ri) by spinning at 4000 × *g* for 10 minutes. The supernatant was discarded and the cell pellets were re-suspended in refrigerated 100 mM calcium chloride solution; first 1 mL CaCl_2_ was used to re-suspend the pellet by gentle pipetting, then the cell suspension from the 5 mL culutre volumes were transferred to pre-chilled 1.5 mL microfuge tubes. Meanwhile, a further 20 mL cold 100 mM CaCl_2_ solution was added to each of the 50 mL culture tubes. These cell suspensions were incubated on ice for 20 minutes. Next, the cell suspensions in the 1.5 mL microfuge tubes were pelleted by centrifugation for 1 min at 6000 × *g*, the supernatant was pipetted out and discarded and the cell pellet was re-suspended in 120 µl of cold 100 mM CaCl_2_ with glycerol at 15%. Larger cell suspensions were pelleted by centrifugation at 4000 × *g* for 10 minutes. The supernatant was discarded and the cell pellets were re-suspended in cold 100 mM CaCl_2_ with 15% glycerol. All these cell pellets were pooled into one volume approximately 1/200 of the original culture volume (i.e., 200 mL of cell culture would be concentrated into 1 mL). Regardless of the original culture size, all HSCC cells were split into 20 µL aliquots that were placed in pre-chilled PCR tubes (StarLab, A10402-3700;), which were then stored in a –80°C freezer for at least one night before use.

### Heat shock transformations of *E. coli cells* for colony forming unit counting

All heat shock transformations were performed in a Bio-Rad C1000 Touch Thermal Cycler (Bio-Rad, 1851148) using the following protocol; 20 min at 0°C, 1 min at 42°C, 3 min at 0°C, infinite-hold at 25°C. For **Figure 1d–f**, 20 ng of purified pR6K-mScarI minipreped DNA was transformed, and for **Figure 2b** 1.66 µL from a 10 µL cloning assembly reaction mix was used for transformations. Post heat-shock, the transformed cells were transferred to 200 µL of SOC (SOB with 0.2% glucose) in a 1.5 mL microfuge tube and then incubated in a Eppendorf ThermoMixer C (Eppendorf, 5382000031) set to 37°C (**Figure 1d–f**) or 30°C (**Figure 2b**) for 1 hour, shaking at 900 rpm. Post recovery, the cultures were serial diluted through seven 10-fold steps, and 5 µL of each dilution spotted onto selective (Kan or Cam) and non-selective plates. These plates were incubated overnight at 37°C or 30°C, as appropriate, and were imaged using a Bio-Rad GelDoc Go Gel Imaging System (Bio-Rad, 12009077) for colony forming unit (CFU) counting the following morning. CFUs were counted from images taken of the plates (example in **Supplementary Figure 3**) and the following formula used to determine transformation efficiency:

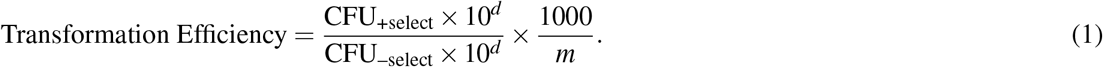

Here, *d* is the dilution factor at which colonies are counted, and *m* is the DNA mass in ng units, reporting transformation efficiency in units of CFU per recipient cell per µg of transformed DNA. For **Figure 2b**, efficiency calculations are not normalized to a specific DNA quantity as this is not known, therefore transformation efficiency is reported as CFU per recipient cell.

### Construction of SHARK strains with *λ*RED

A custom *λ*RED plasmid was created, called pREMORA, by placing the *β, γ* and exo operon under the regulation of a salicylic acid inducible promoter on a temperature sensitive plasmid, pSC101TS that also contains a beta-lactamase gene for antibiotic selection (*ampR*). A *recA* gene with a unregulated P_*lexA*_ promoter was also placed on pREMORA, enabling *in vivo* recombination in *recA*^−^ strains, such as DH10B. The *λ*RED operon and *recA* were taken from pREDCas9, a gift from Tao Chen (Addgene plasmid No.71541) [40]; the salicylic acid inducible promoter and transcription factor were from pAJM.771, a gift from Christopher Voigt (Addgene plasmid No.108534) [11], and the pSC101TS backbone was from pE-FLP, a gift from from Drew Endy and Keith Shearwin (Addgene plasmid No. 45978) [4]. pREMORA was constructed using Golden-gate cloning, and was assembled in two steps; first the *λ*RED operon and P_*lexA*_-*recA* gene were placed under PSalTC regulation on pAJM.771, this plasmid has a p15A-Kanamycin backbone. Then, the origin and selection marker were changed to that of pSC101TS-*ampR* for easy curing post recombination. Each SHARK design was first cloned onto a replicative pBBR1 plasmid with a FRT flanked SpecR resistance marker. This vector was created from various components and the full plasmid sequence can be found in **Supplementary Data 1**. The *λ*CI transcription unit and SHARK1-Pir transcription unit were first cloned individually into p15A-Kan vectors and then these were combined onto one pBBR1-SpecR-FRT vector in a second round of Golden-Gate assembly. All the parts for the *λ*CI and Pir genes were taken from the CIDAR MoClo Extension Volume I kit, a gift from Richard Murray (Addgene kit No. 1000000161), except the wildtype Pir promoter and RBS, which were ordered as oligos and annealed to use for the Golden-Gate assembly. For SHARK2–7, the promoters and RBSs were modified using Golden-Gate, with oligo annealed parts for P_*tet*_ and B0033, and a PCR amplified fragment was used for RiboJ-B0064. The CDSs were modified using PCR mediated site-directed mutagenesis and blunt-end ligation. The SHARK8–10 designs were created by PCR mediated site-directed mutagenesis of SHARK2 to swap the P_*tet*_ promoter to a weak constitutive promoter (P_J23117_, P_J23114_ and P_J23103_). Fully annotated plasmid sequences can be found as **Supplementary Data 1**.

For integrating the SHARK cassettes into the *E. coli* Marionette-Clo genome, cells transformed with pREMORA were made HSCC using the protocol described earlier; however, with 2 key changes; the cells were incubated at 30°C as the plasmid is temperature sensitive, and the optical density at 700 nm (OD700) was monitored so that the culture could be induced with 100 µM salicylate at and OD700 of 0.5 to activate the *λ*RED operon. The culture was then further incubated shaking at 30°C for 30 minutes before proceeding to the incubation on ice step of the protocol.

All SHARK cassettes were PCR amplified using Q5 High-Fidelity 2X Master Mix (New England Biolabs, M0492) in 50 µL reactions with 10 ng of template plasmid DNA. PCRs was performed in a C1000 Touch Thermal Cycler (Bio-Rad, 1851148) for 25 cycles. The DNA template was digested away using Dpni (New England Biolabs, R0176) by adding 1 µL of the enzyme directly to the reaction mix and then incubating at 37°C for 1 hour. The reaction mix was then run on an agarose gel by electrophoresis, and the DNA band extracted using a Monarch Spin DNA Gel Extraction Kit (New England Biolabs, T1120), following manufacturer’s guidelines. After purification, 200 ng of linear DNA was used for heat shock transformation into the *λ*RED competent Marionette-Clo cells. Post transformation, the cells were recovered at 37°C and were plated on spectinomycin selection plates and incubated overnight at 37°C. The following day, 6–8 single colonies for each SHARK design were passaged onto new spectinomycin selection plates to ensure single clone isolation and further dilution of pREMORA from the population. The following day, colony PCR was conducted to ensure genome integration of the cassette, then a single colony for each SHARK strain was inoculated into 2 mL SOB with spectinomycin and cultured overnight at 37°C shaking at 250 rpm. The following day, the SHARK strain was made chemical competent in a 5 mL culture volume and the cells transformed with 50 ng of miniprepped pE-FLP DNA. Cells were then incubated at 30°C to maintain the temperature sensitive plasmid, and carbenicillin used for selection. The following day, a single colony for each SHARK strain was passaged on a new carbenicillin plate to ensure curing of the selection marker from the genome. The following day, a single colony was inoculated into 200 µL of LB media and cultured in a Eppendorf ThermoMixer C (Eppendorf, 5382000031) at 42°C for 2 hours, then 37°C for 6 hours. At the end of the day, the culture was streaked out onto non-selective LB agar plates. The following day, a single colony was picked to be the final SHARK strain used in all subsequent experiments. A final colony PCR was conducted to ensure successful curing of the selection marker (see **Supplementary Figure 1b**).

### Cloning methods

PCR amplicons for cloning purposes and *λ*RED were amplified using Q5 High-Fidelity 2X Master Mix (New England Biolabs, M0492) and run on agarose gels for DNA band separation and extractions using a Monarch Spin DNA Gel Extraction Kit (New England Biolabs, T1120). Colony PCRs were done using OneTaq Quick-Load 2X Master Mix (New England Biolabs, M0486). Plasmids were built using either Golden-Gate assembly, or blunt-end ligation (BEL) for site directed mutagenesis. BsaI-HFv2 (New England Biolabs, R3733S, 20,000 units/mL) and T4 DNA ligase (New England Biolabs, M0202S, 400,000 units/mL) were used for all Golden-Gate reaction. Each reaction was set to a total volume of 10 µL containing 40 fmol of DNA for the recipient backbone, 80 fmol of DNA for each insert, 1 µL BsaI-HFv2, 1 µL T4 DNA ligase, 1 µL T4 DNA ligase buffer, and water. The Golden-Gate reactions were incubated for 1 hour at 37°C in a standard incubator. For BEL cloning reactions, each reaction was set to a total volume of 10 µL containing 200 ng of PCR amplified linear DNA, 1 µL T4 DNA ligase, 1 µL T4 polynucleotide kinase (New England Biolabs, M0201L, 10,000 units/mL), 1 µL T4 DNA ligase buffer and water. The BEL reactions were incubated for 1 hour at 37°C in a standard incubator. For regular cloning reactions, 5 µL of the reaction mix was used to transform the chemical competent *E. coli* cloning strain, but only 1.66 µL was used to transform each competent cell aliquot for experiments presented in **Figure 2b**.

### Microplate reader experiments

All microplate reader experiments were conducted in a SpectraMax iD5 Multi-Mode Microplate Reader (Molecular Devices) in 96-well flat-bottom Sterilin Microtitre Plates (STERLIN, 734-0482). For all experiments *E. coli* cells were transformed fresh with the respective plasmids of interest. Untransformed *E. coli* cells were also streaked out, to be used for positive controls for growth and for subtracting cell autofluorescence in the analysis, where relevant. Three colonies from each strain of interest were inoculated into 200 µL of M9-glucose media in 2 mL square-well 96-well deep-well plates (Merck, AXYP2MLSQC); the media contains the relevant antibiotics, but no inducers unless relevant. All deep-well 96-well plates were sealed with a breathable membrane (Starlab E2796-3015) and incubated at 37°C in a plate shaker (Stuart SI505 Microtitre Plate Shaker Incubator) set to 750 rpm for 16–18 hours overnight. The following morning, the cultures were diluted 200-fold by transferring 1 µL from the overnight wells to 200 µL of fresh media of the same conditions, also set up in deep-well plates. These diluted cultures (referred to as precultures) were also incubated at 37°C in a plate shaker set to 750 rpm, but for only 3 hours. After 3 hours, the precultures were diluted 400-fold by transferring 0.5 µL to 200 µL of fresh media in a 96-well flat-bottom Sterilin Microtitre Plates (STERLIN, 734-0482); this media contained the relevant inducer conditions. For every plate-reader experiment three “blank” wells were set up of M9-glucose media with no inoculant. The microplate was then loaded into a SpectraMax iD5 Multi-Mode Microplate Reader (Molecular Devices). The plate was set to take the following measurements every 10 minutes for a total of 15 hours; absorbance at 700 nm (OD700), green fluorescence (excitation at 485, emmision at 515 nm, with a sensitivity gain of 500 a.u.) and red fluorescence (excitation at 565, emmision at 610 nm, with a sensitivity gain of 750 a.u.).

### Microplate reader data analysis for plasmid copy number estimation and sfGFP repression

All plate reader data was exported as a text file from the SpectraMax iD5 instrument software and converted to a CSV format for analysis using Python. Raw data was preprocessed by first subtracting the background absorbance and autofluorescence from the designated “blank” wells. Then a “hard” lower-bound threshold was set for all OD700 values (i.e., all OD700 measurements that were below the threshold were set to the threshold value). This was done because the cell density for the first few hours of the experiment was below the detection limit of the plate reader, generating a lot of noise, and subtracting the blanks caused some negative OD700 values which are problematic for growth rate analysis. Setting a hard lower-bound threshold mitigated these issues for subsequent rate analysis steps. For each experiment, the lower-bound threshold was set to a twenty fifth of the value of the maximum OD700 measured for the designated positive growth control wells. The fluorescence data did not require thresholding like OD700. The absorbance and fluorescence values were further processed by running a smoothing function (6-window moving average for each well) to reduce noise and occurrence of sharp peaks in the subsequent analysis steps. Cell growth rate (hr^−1^) and fluorescence production rate per cell (OD700) were calculated for each well using:

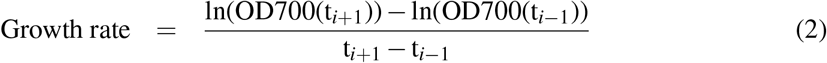

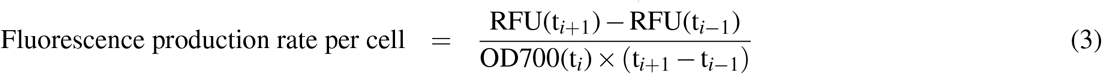

Fluorescence production rate at maximum growth rate was used to estimate plasmid copy number **(Supplementary Figure 4)** and report sfGFP expression **(Supplementary Figure 5)**. Plasmid copy number was estimated by dividing mScarlet-I expression from cells carrying pR6K-mScarI by cells carrying p15AmScarI, then multiplying that value by 9 [32–34] to get estimated plasmid copy number per genome. sfGFP fluorescence expression reported in **Figure 1e** is normalized to non-fluorescent cells. Representative data for growth and fluorescence curves are presented in **Supplementary Figure 7**.

### Computational tools

Microsoft Excel 2025 was used to tabulate and calculate all reported values for CFUs and transformation efficiencies. All microplate reader analysis and graphing of figures was done using custom scripts in Python, version 3.9.7, using the Jupyter Lab interface, version 3.2.1. All diagrams and figure panels were assembled using Inkscape version 1.4.2.

### Availability of plasmids and strains

The following plasmids and strains are in the process of being deposited at Addgene: pR6K-mScarI, pPRO1O2-GFP, pSHARK2, pSHARK6, pSHARK7, pSHARK8, pREMORAv1, SHARK2, SHARK6, SHARK7, SHARK8.

## DATA AVAILABILITY

**Supplementary Data 1** contains sequences of all key genetic parts in FASTA format and annotated plasmid sequences for all SHARK designs in GenBank format.

## ACKNOWLEDGMENTS

S.N.-H.J. was supported by a Dstl-funded PhD Studentship. T.E.G. was additionally supported by the UKRI Engineering Biology Mission Award CYBER under BBSRC grant BB/Y007638/1, and a Royal Society University Research Fellowship grant URF*\*R*\*221008. The funders had no role in study design, data collection and analysis, and decision to publish or preparation of the manuscript.

## AUTHOR CONTRIBUTIONS

S.H.-N.J. conceived the study, built the strains, carried out all experiments, performed the analysis, and wrote the initial draft of the manuscript. T.E.G. secured funding and aided in the analysis. T.E.G., C.J. and D.U. supervised the work and edited of the final manuscript.

