## Supplementary Information for "SHARK: a specialized host for assembling R6K plasmids"

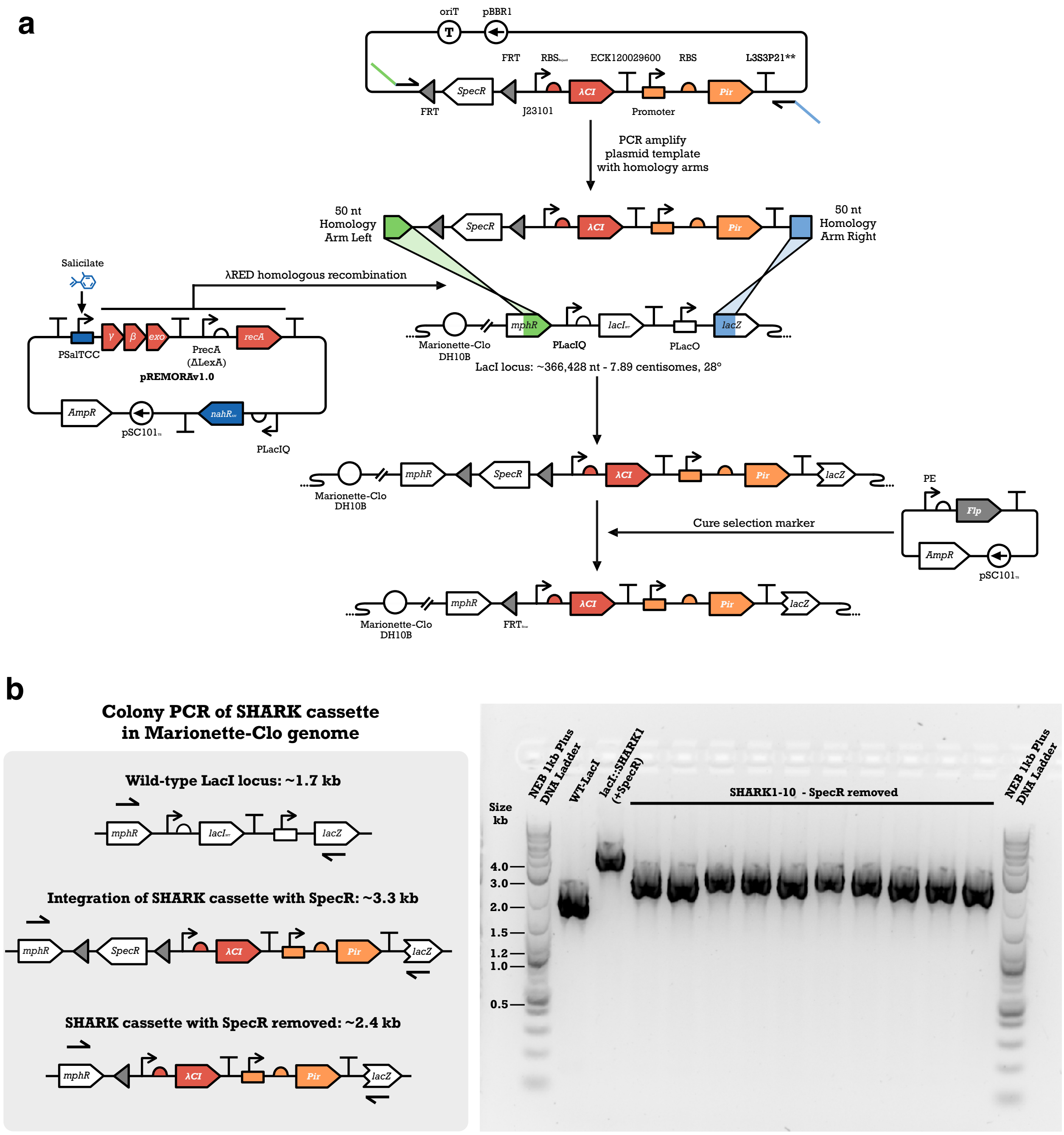

**Supplementary Figure 1: Construction of SHARK strains using $\lambda$-RED recombineering.** (**a**) All SHARK cassette designs were first constructed on a pBBR1 plasmid with a spectinomycin resistance marker (adenylyltransferase aadA, SpecR) flanked by flippase recognition target (FRT) sequences. These plasmids were used as the templates to PCR amplify the SHARK cassette. The overhangs of the PCR primers include 50 base-pairs of homology downstream of nucleotide 910 of mphR (left) and nucleotide 162 of lacZ (right) coding sequence. All PCR products were gel purified and approximately 200 ng of DNA was transformed to chemical competent $\lambda$RED-ready cells. Successful integration would replace the wild-type LacI gene, as well as remove the PlacO promoter and first 162 nt of lacZ. Stable integrants were selected for using spectinomycin post transformation, and passaging once on selective solid media, and once in selective liquid media. Final candidate clones were then made chemical competent and transformed with a Flippase (FLP) expressing plasmid with a temperature sensitive origin of replication (pSC101-TS). Single colonies would be propagated on carbenicillin media for at least one passage, then the plasmid would be cured by growing cells in liquid cultures for 2 hours at 42°C, followed by 4–6 hours at 37°C, inhibiting pSC101-TS replication. (**b**) All SHARK strains were verified for successful integration using colony PCR. The expected sizes of the negative control amplicon (wild-type LacI), intermediate amplicon (with SpecR marker), and the final SHARK cassettes (SpecR marker cured) are approximately 1.7 kb, 3.3 kb and 2.4 kb, respectively. The colony PCR demonstrates the expected results for successful genome integration and curing of the SpecR marker using our described $\lambda$RED recombineering method.

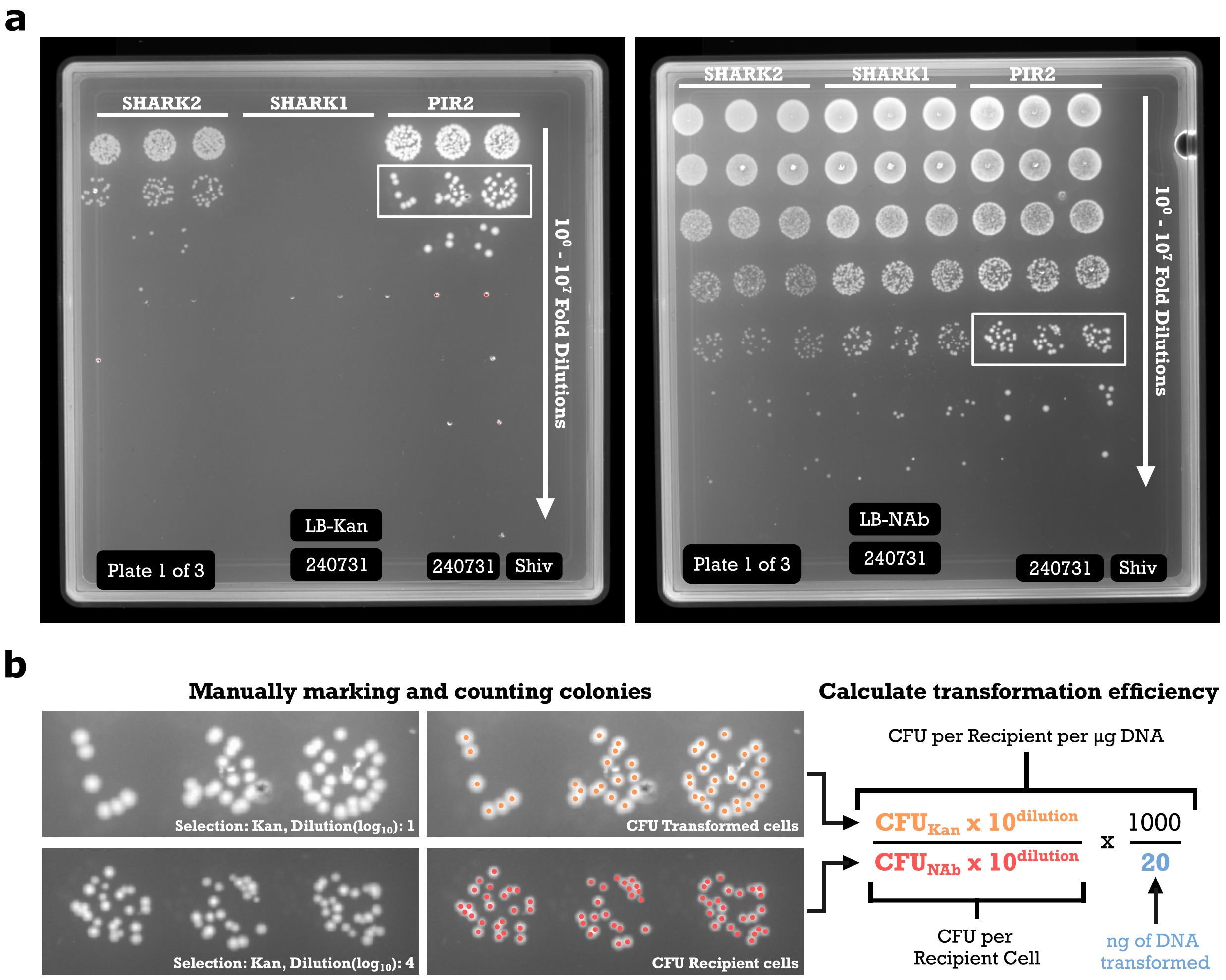

**Supplementary Figure 2: Sample of transformation efficiency characterization experiment.** (**a**) PIR2, SHARK1 and SHARK2 transformations with pR6K-mScarI were serially diluted and spotted on square plates with and without kanamycin selection. White rectangles show examples of cell spots that were counted. Hand written labels on the plates are overlaid with digital counterparts (white text in black-filled boxes) for clarity. (**b**) Zoomed sections from panel (a) with manually counted colonies highlighted in orange and red. These CFU values and respective dilution were used to calculate transformation efficiency.

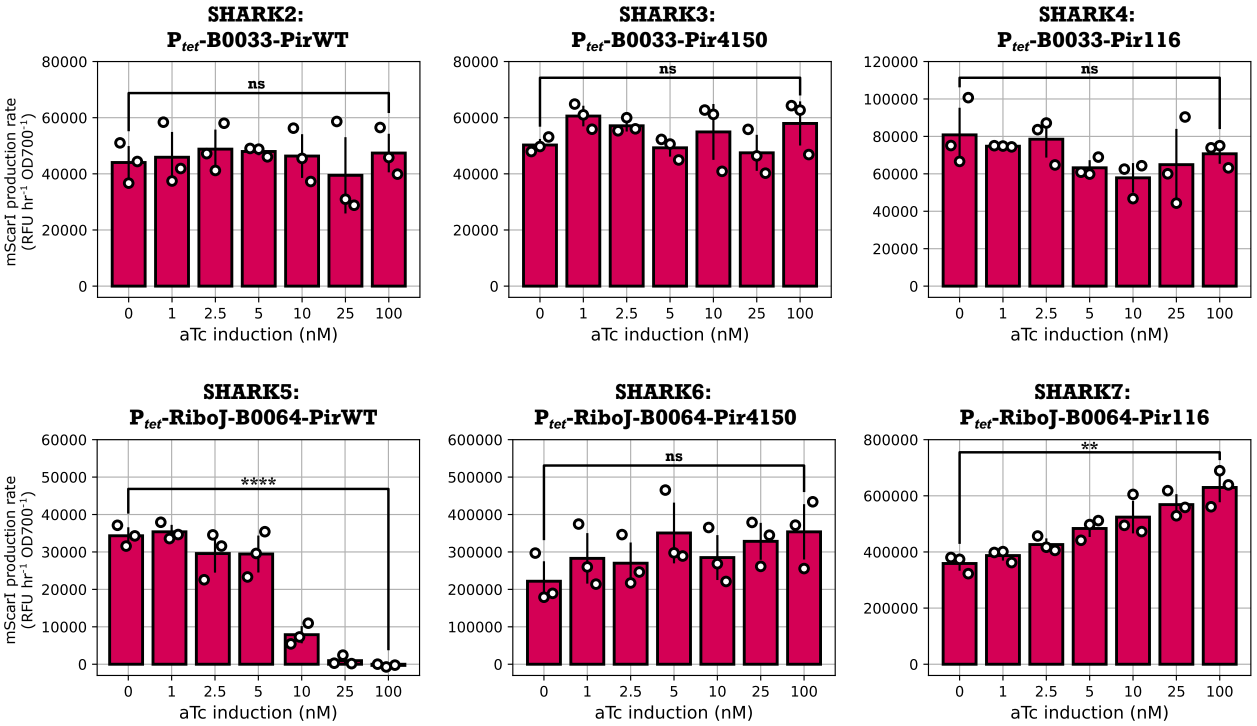

**Supplementary Figure 3: Control of RK6 plasmid copy number in SHARK strains.** SHARK2–7 have their Pir gene expressed from an aTc inducible $P_{tet}$ promoter. To test if stronger expression of the Pir gene affected R6K plasmid copy number, we used mScarlet-I constitutively expressed from the pR6K-mSarI plasmid as an indicator for plasmid abundance and varied aTc concentration. Higher production rates of mScarlet-I denote a higher plasmid copy number. ns: no statistically significant difference (Student's $t$-test, $p > 0.05$); **: statistically significant difference (Student's $t$-test, $p \leq 0.05$); ****: statistically significant difference (Student's $t$-test, $p \leq 0.0001$).

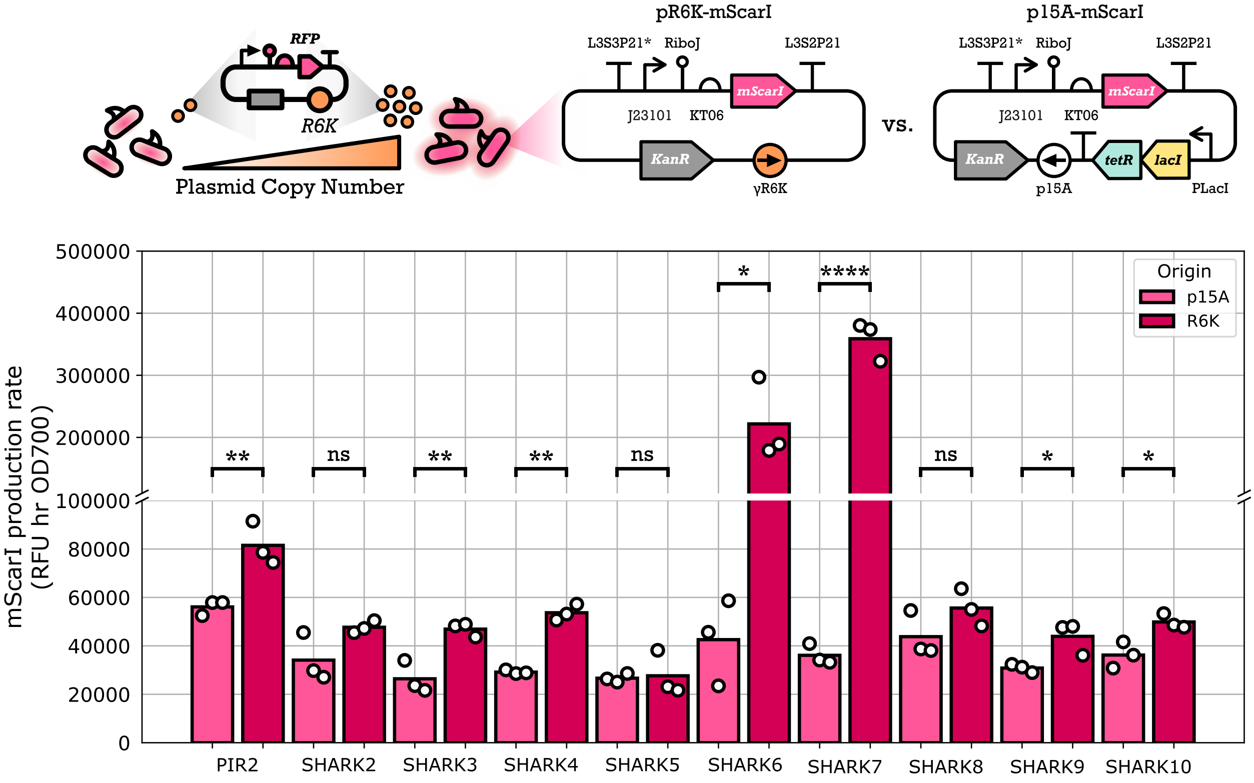

**Supplementary Figure 4: mScarlet-I production rates used to estimate plasmid copy number.** PIR2 and the SHARK strains were transformed with pR6K-mScarI and p15A-mScarI. The growth and fluorescence of these strains were measured in a kinetic microplate reader experiment. The mScarlet-I fluorescence production rate was calculated for all strains and the value at maximum growth rate is reported here for each samples. For 7 of the 10 strains shown, the difference in mScarlet-I fluorescence production rate between R6K and p15A plasmids was significant (Student T-test). Regardless, plasmid copy number was estimated by normalizing mScarlet-I production rate of R6K plasmids by p15A for each strain, then multiplying by 9 to get PCN, these are reported in **Figure 1d**. ns: no statistically significant difference (Student's $t$-test, $p > 0.05$); *: statistically significant difference (Student's $t$-test, $p \leq 0.05$); **: statistically significant difference (Student's $t$-test, $p \leq 0.01$; ****: statistically significant difference (Student's $t$-test, $p \leq 0.0001$.

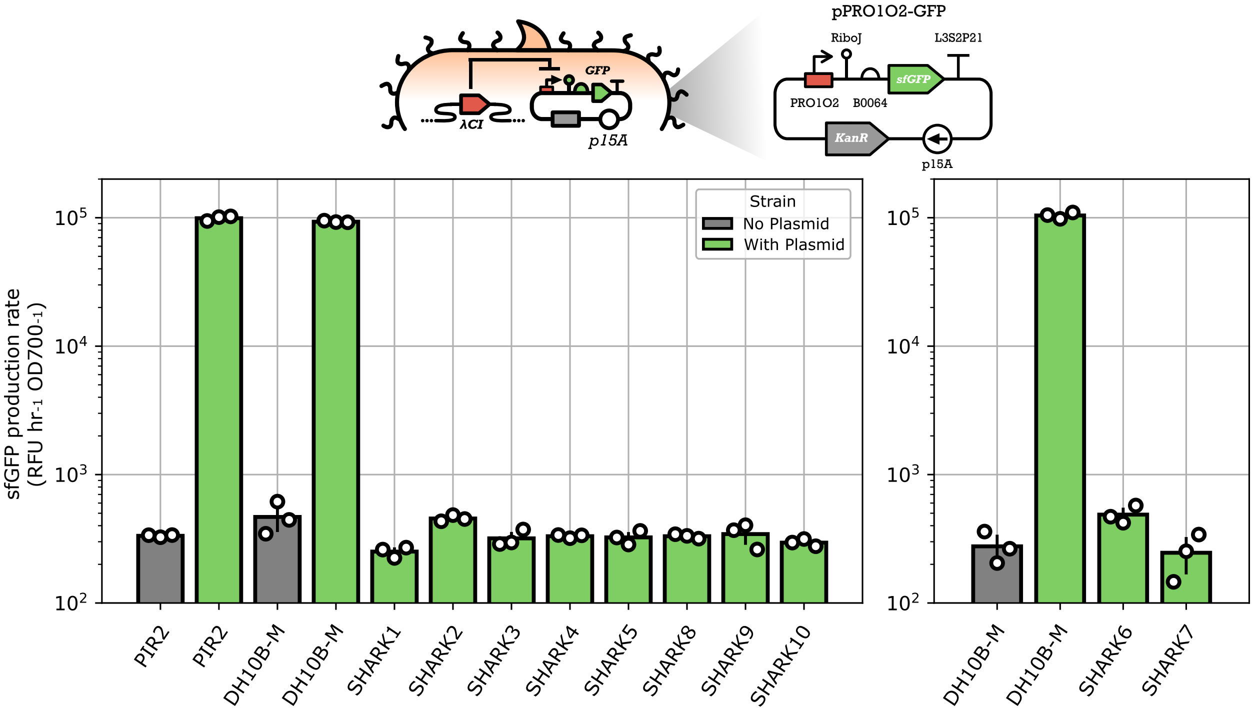

**Supplementary Figure 5: sfGFP production rates for pPRO1O2-sfGFP expression in SHARK.** PIR2 and the SHARK strains were transformed with pPRO1O2-sfGFP, and their growth and fluorescence was measured in a kinetic microplate reader experiment. The sfGFP fluorescence production rate was calculated for all strains and the value at each samples maximum growth rate is reported here. These production rates were normalized to the respective untransformed cell autofluorescence control, and this is reported in **Figure 1e**. sfGFP experiments for SHARK6 and SHARK7 were done in a different run to the rest of the SHARK strains, hence the data being represented in two plots with their own respective controls.

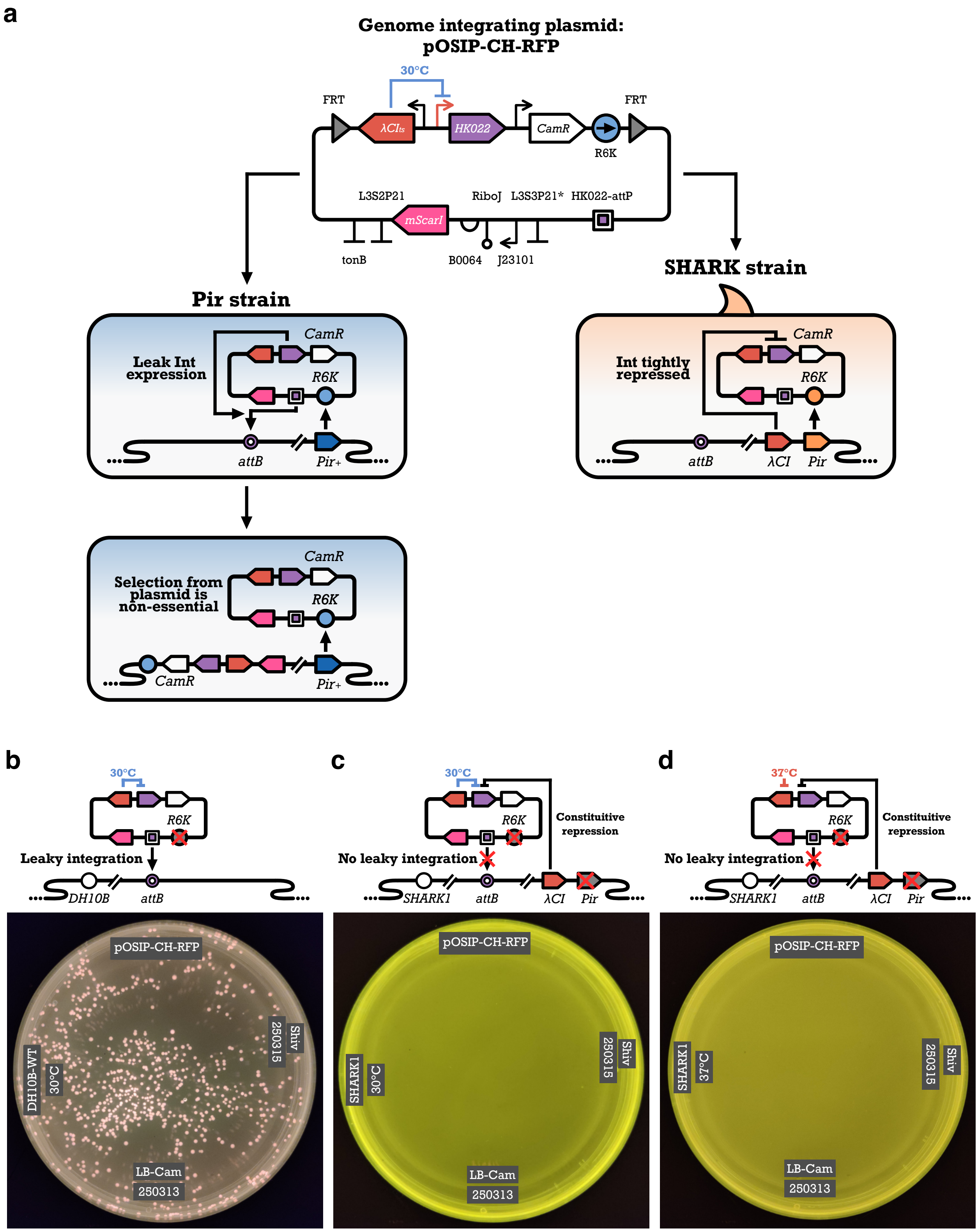

**Supplementary Figure 6: Genome integration of pOSIP plasmid without activation.** (**a**) pOSIP plasmids are designed for efficient *E. coli* genome engineering. The backbone of these plasmids have a R6K origin and a temperature sensitive λCI (λ$CI_{ts}$) which inhibits a phage integrase (i.e., HK022) that is expressed in the cloning strain. However, leaky expression of the integrase can result in unintended genome integration of the plasmid, conferring antibiotic resistance to the host genome and making plasmid maintenance unstable after selection. To mitigate this issue, SHARK strains have an additional $\lambda$CI gene to ensure tight repression of the integrase. (**b**) To demonstrate leaky expression of the integrase, chemical competent DH10B cells without any Pir gene were transformed with an RFP expressing pOSIP plasmid (pOSIP-CH-RFP) and were kept at 30°C to inhibit expression of the integrase. However, hundreds of colonies continued to grow on selective plates, showing that repression of the temperature sensitive $\lambda$CI is insufficient to prevent genome integration. The pOSIP-CH-RFP plasmid was transformed to SHARK1, which does not have a functional Pir cassette, but does have tight repression of the $\lambda$-promoters. SHARK1 transformed with pOSIP-CH-RFP were kept at (**c**) 30°C, as well as (**d**) 37°C to turn off the temperature sensitive $\lambda$CI protein from the plasmid. We found no colony growth from these transformations. Hand written labels on the plate images are overlaid with digital counterparts for clarity. Note, the yellow-green colour of the plates in (**c**) and (**d**) is from the camera auto-adjusting the colour correction and is detecting the autofluorescence from the LB agar plate.

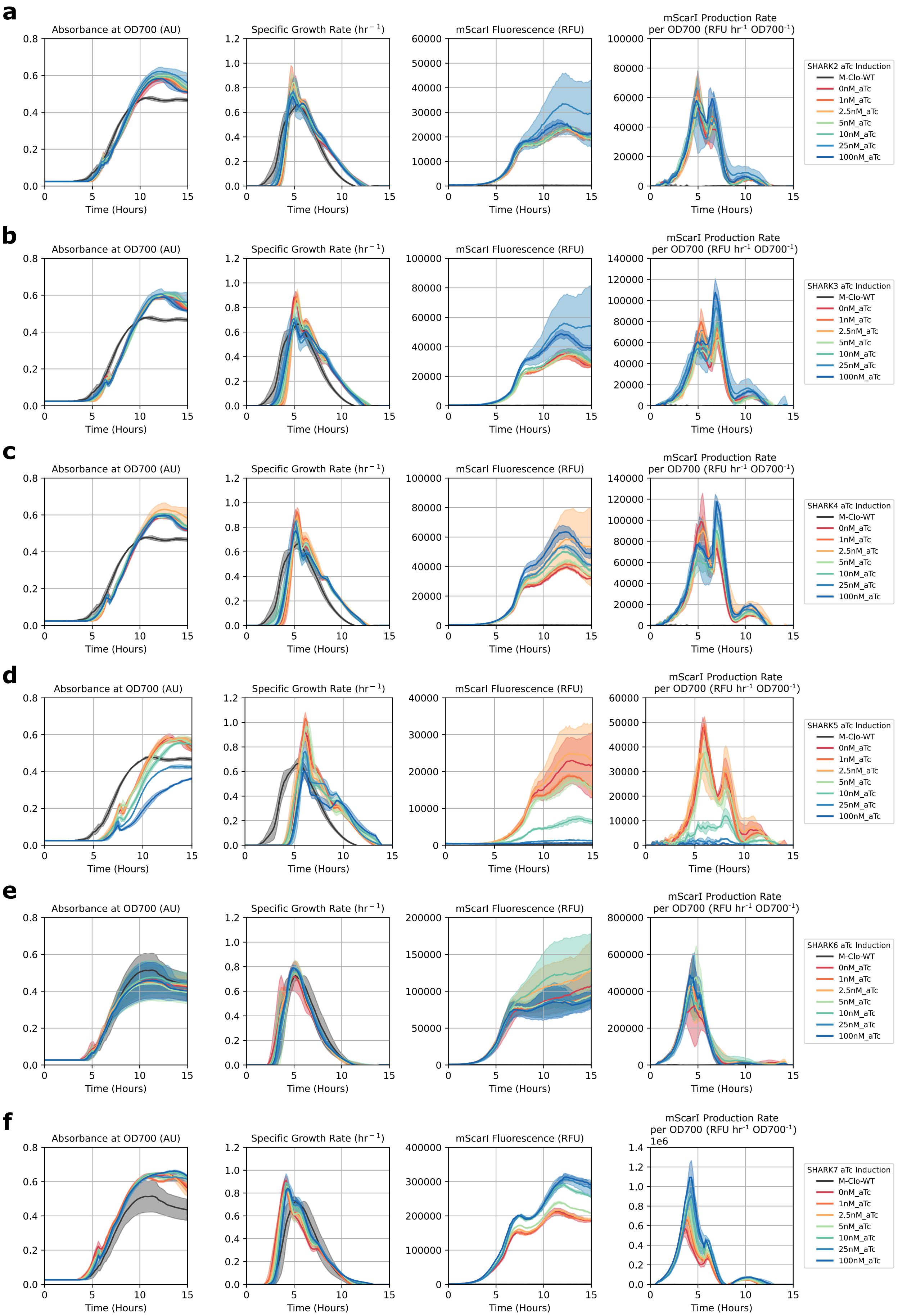

(Figure continued on next page...)

g

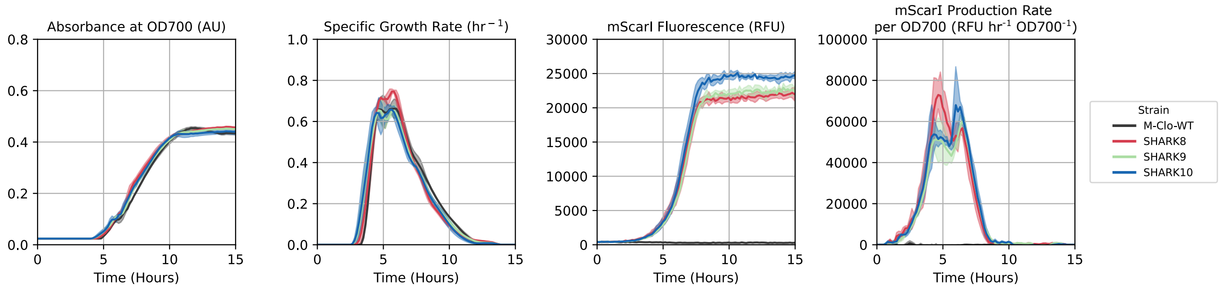

**Supplementary Figure 7: Time course measurements of SHARK strains with pR6K-mScarI.** Time course measurements for Marionette-Clo (M-Clo-WT) and (**a**) SHARK2, (**b**) SHARK3, (**c**) SHARK4, (**d**) SHARK5, (**e**) SHARK6, and (**f**) SHARK7 cultured in varying concentrations of aTc. (**g**) Time course measurements for Marionette-Clo (M-Clo-WT) and SHARK8–10 strains.

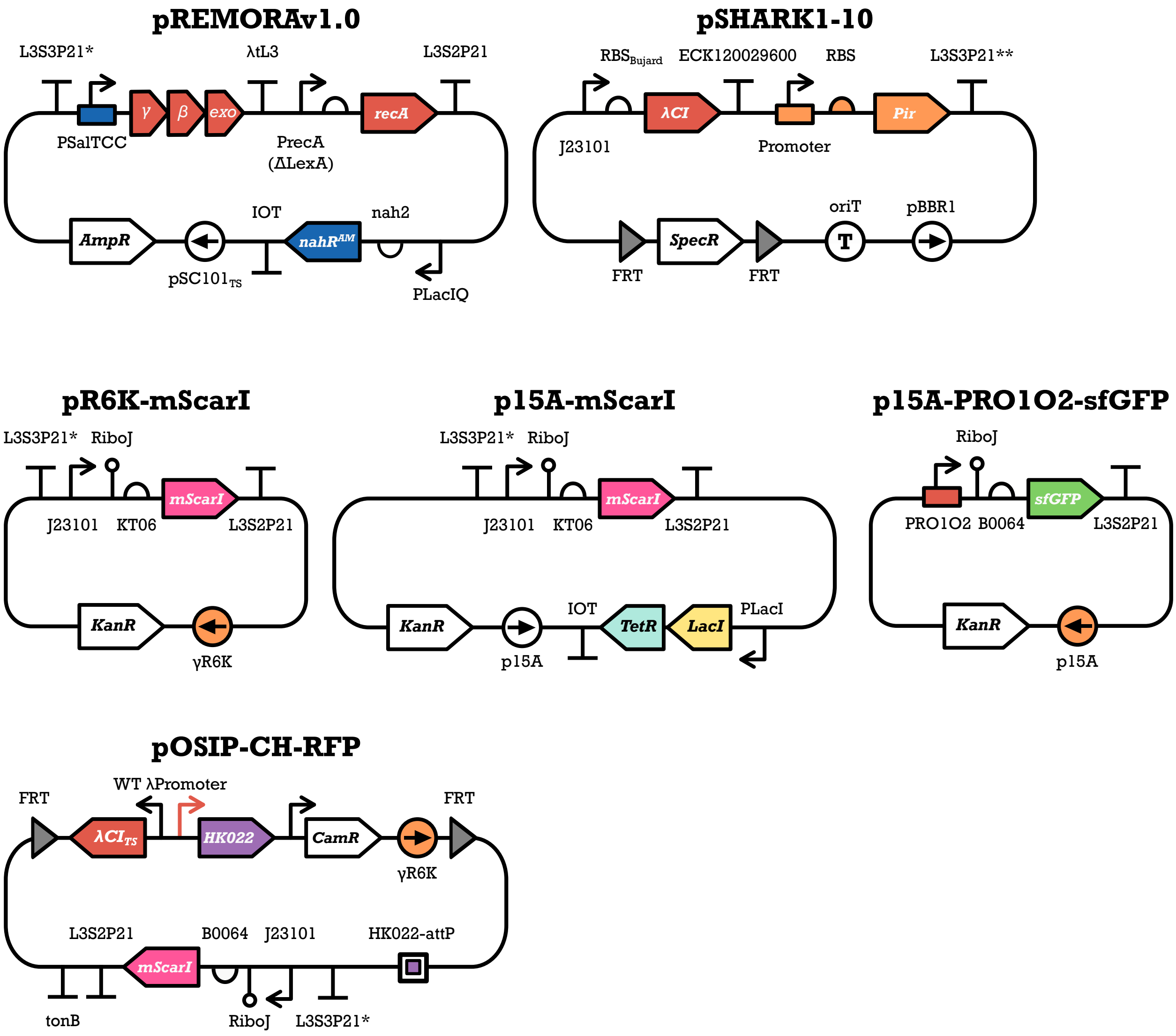

**Supplementary Figure 8: Plasmid maps.** For pSHARK1–10, regulatory and Pir protein coding sequence combinations are shown in **Table 1**.
