## Supplementary Data 1 for "SHARK: a specialized host for assembling R6K plasmids": dna_parts.fasta

>Wildtype_Pir-UTr
TTAAACATGAGTGGATAGTACGTTGCTAAAACATGAGATAAAAATTGACTCTCATGTTATTGGCGTTAAGATATACAGAATGATGAGGTTTTTTT

>PJ23101
TTTACAGCTAGCTCAGTCCTAGGTATTATGCTAGC

>Ptet
TCCCTATCAGTGATAGAGATTGACATCCCTATCAGTGATAGAGATACTGAGCAC

>PJ23117
TTGACAGCTAGCTCAGTCCTAGGGATTGTGCTAGC

>PJ23114
TTTATGGCTAGCTCAGTCCTAGGTACAATGCTAGC

>PJ23103
CTGATAGCTAGCTCAGTCCTAGGGATTATGCTAGC

>PRO1O2
TGCTGTTCCGCTGGGCATGCTGAGCTAACACCGTGCGTGTTGACAATTTTACCTCTGGCGGTGATAATGGTTGCAGC

>RiboJ
AGCTGTCACCGGATGTGCTTTCCGGTCTGATGAGTCCGTGAGGACGAAACAGCCTCTACAAATAATTTTGTTTAA

>B0033
TCTAGAGTCACACAGGACTACTAGA

>B0064
TACTAGAGAAAGAGGGGAAATACTAGA

>RBSBujard
GAATTCATTAAAGAGGAGAAAGGTACCA

>KT06
CATAATATTTCTCTTCCGGTATAAA

>Lambda-CI
ATGAGCACAAAAAAGAAACCATTAACACAAGAGCAGCTTGAGGACGCACGTCGCCTTAAAGCAATTTATGAAAAAAAGAAAAATGAACTTGGCTTATCCCAGGAATCTGTCGCAGACAAGATGGGGATGGGGCAGTCAGGCGTTGGTGCTTTATTTAATGGCATCAATGCATTAAATGCTTATAACGCCGCATTGCTTGCAAAAATTCTCAAAGTTAGCGTTGAAGAATTTAGCCCTTCAATCGCCAGAGAAATCTACGAGATGTATGAAGCGGTTAGTATGCAGCCGTCACTTAGAAGTGAGTATGAGTACCCTGTTTTTTCTCATGTTCAGGCAGGGATGTTCTCACCTGAGCTTAGAACCTTTACCAAAGGTGATGCGGAGAGATGGGTAAGCACAACCAAAAAAGCCAGTGATTCTGCATTCTGGCTTGAGGTTGAAGGTAATTCCATGACCGCACCAACAGGCTCCAAGCCAAGCTTTCCTGACGGAATGTTAATTCTCGTTGACCCTGAGCAGGCTGTTGAGCCAGGTGATTTCTGCATAGCCAGACTTGGGGGTGATGAGTTTACCTTCAAGAAACTGATCAGGGATAGCGGTCAGGTGTTTTTACAACCACTAAACCCACAGTACCCAATGATCCCATGCAATGAGAGTTGTTCCGTTGTGGGGAAAGTTATCGCTAGTCAGTGGCCTGAAGAGACGTTTGGCTGA

>Pir
ATGAGACTCAAGGTCATGATGGACGTGAACAAAAAAACGAAAATTCGCCACCGAAACGAGCTAAATCACACCCTGGCTCAACTTCCTTTGCCCGCAAAGCGAGTGATGTATATGGCGCTTGCTCCCATTGATAGCAAAGAACCTCTTGAACGAGGGCGAGTTTTCAAAATTAGGGCTGAAGATCTTGCAGCGCTCGCCAAAATCACCCCATCGCTTGCTTATCGACAATTAAAAGAGGGTGGTAAATTACTTGGTGCCAGCAAAATTTCGCTAAGAGGGGATGATATCATTGCTTTAGCTAAAGAGCTTAACCTGCCCTTTACTGCTAAAAACTCCCCTGAAGAGTTAGATCTTAACATTATTGAGTGGATAGCTTATTCAAATGATGAAGGATACTTGTCTTTAAAATTCACCAGAACCATAGAACCATATATCTCTAGCCTTATTGGGAAAAAAAATAAATTCACAACGCAATTGTTAACGGCAAGCTTACGCTTAAGTAGCCAGTATTCATCTTCTCTTTATCAACTTATCAGGAAGCATTACTCTAATTTTAAGAAGAAAAATTATTTTATTATTTCCGTTGATGAGTTAAAGGAAGAGTTAATAGCTTATACTTTTGATAAAGATGGAAATATTGAGTACAAATACCCTGACTTTCCTATTTTTAAAAGGGATGTGTTAAATAAAGCCATTGCTGAAATTAAAAAGAAAACAGAAATATCGTTTGTTGGCTTCACTGTTCATGAAAAAGAAGGAAGAAAAATTAGTAAGCTGAAGTTCGAATTTGTCGTTGATGAAGATGAATTTTCTGGCGATAAAGATGATGAAGCTTTTTTTATGAATTTATCTGAAGCTGATGCAGCTTTTCTCAAGGTATTTGATGAAACCGTACCTCCCAAAAAAGCTAAGGGGTGA

>Pir4150
ATGAGACTCAAGGTCATGATGGACGTGAACAAAAAAACGAAAATTCGCCACCGAAACGAGCTAAATCACACCCTGGCTCAACTTCCTTTGCCCGCAAAGCGAGTGATGTATATGGCGCTTGCTCCCATTGATAGCAAAGAACCTCTTGAACGAGGGCGAGTTTTCAAAATTAGGGCTGAAGATCTTGCAGCGCTCGCCAAAATCACCCCATCGCTTGCTTATCGACAATTAAAAGAGGGTGGTAAATTACTTGGTGCCAGCAAAATTTCGCTAAGAGGGGATGATATCATTGCTTTAGCTAAAGAGCTTAACCTGCCCTTTATTGCTAAAAACTCCTCTGAAGAGTTAGATCTTAACATTATTGAGTGGATAGCTTATTCAAATGATGAAGGATACTTGTCTTTAAAATTCACCAGAACCATAGAACCATATATCTCTAGCCTTATTGGGAAAAAAAATAAATTCACAACGCAATTGTTAACGGCAAGCTTACGCTTAAGTAGCCAGTATTCATCTTCTCTTTATCAACTTATCAGGAAGCATTACTCTAATTTTAAGAAGAAAAATTATTTTATTATTTCCGTTGATGAGTTAAAGGAAGAGTTAATAGCTTATACTTTTGATAAAGATGGAAATATTGAGTACAAATACCCTGACTTTCCTATTTTTAAAAGGGATGTGTTAAATAAAGCCATTGCTGAAATTAAAAAGAAAACAGAAATATCGTTTGTTGGCTTCACTGTTCATGAAAAAGAAGGAAGAAAAATTAGTAAGCTGAAGTTCGAATTTGTCGTTGATGAAGATGAATTTTCTGGCGATAAAGATGATGAAGCTTTTTTTATGAATTTATCTGAAGCTGATGCAGCTTTTCTCAAGGTATTTGATGAAACCGTACCTCCCAAAAAAGCTAAGGGGTGA

>Pir116
ATGAGACTCAAGGTCATGATGGACGTGAACAAAAAAACGAAAATTCGCCACCGAAACGAGCTAAATCACACCCTGGCTCAACTTCCTTTGCCCGCAAAGCGAGTGATGTATATGGCGCTTGCTCCCATTGATAGCAAAGAACCTCTTGAACGAGGGCGAGTTTTCAAAATTAGGGCTGAAGATCTTGCAGCGCTCGCCAAAATCACCCCATCGCTTGCTTATCGACAATTAAAAGAGGGTGGTAAATTACTTGGTGCCAGCAAAATTTCGCTAAGAGGGGATGATATCATTGCTTTAGCTAAAGAGCTTAACCTGCTCTTTACTGCTAAAAACTCCCCTGAAGAGTTAGATCTTAACATTATTGAGTGGATAGCTTATTCAAATGATGAAGGATACTTGTCTTTAAAATTCACCAGAACCATAGAACCATATATCTCTAGCCTTATTGGGAAAAAAAATAAATTCACAACGCAATTGTTAACGGCAAGCTTACGCTTAAGTAGCCAGTATTCATCTTCTCTTTATCAACTTATCAGGAAGCATTACTCTAATTTTAAGAAGAAAAATTATTTTATTATTTCCGTTGATGAGTTAAAGGAAGAGTTAATAGCTTATACTTTTGATAAAGATGGAAATATTGAGTACAAATACCCTGACTTTCCTATTTTTAAAAGGGATGTGTTAAATAAAGCCATTGCTGAAATTAAAAAGAAAACAGAAATATCGTTTGTTGGCTTCACTGTTCATGAAAAAGAAGGAAGAAAAATTAGTAAGCTGAAGTTCGAATTTGTCGTTGATGAAGATGAATTTTCTGGCGATAAAGATGATGAAGCTTTTTTTATGAATTTATCTGAAGCTGATGCAGCTTTTCTCAAGGTATTTGATGAAACCGTACCTCCCAAAAAAGCTAAGGGGTGA

>mScarlet-I
ATGGTCAGTAAAGGCGAAGCAGTTATCAAAGAGTTCATGCGCTTCAAAGTTCATATGGAAGGGTCGATGAACGGGCACGAATTTGAAATTGAAGGCGAAGGCGAAGGCCGCCCATATGAAGGGACCCAAACCGCAAAGCTTAAGGTTACTAAAGGCGGTCCATTACCCTTTTCGTGGGACATTTTAAGCCCACAGTTTATGTACGGGAGTCGCGCTTTCATCAAGCACCCTGCGGACATCCCAGATTACTACAAACAGTCTTTCCCCGAGGGGTTCAAGTGGGAGCGCGTGATGAACTTCGAGGATGGCGGAGCCGTGACGGTCACCCAAGATACCTCTTTGGAGGACGGTACGTTGATCTACAAAGTGAAATTGCGTGGCACGAATTTTCCACCTGATGGGCCTGTCATGCAGAAAAAGACAATGGGATGGGAAGCTTCCACGGAGCGCCTTTACCCAGAGGACGGTGTTCTTAAAGGGGATATCAAAATGGCGCTGCGTCTTAAAGATGGAGGCCGCTACCTGGCGGACTTCAAGACTACTTACAAGGCCAAAAAACCAGTGCAGATGCCGGGTGCGTACAATGTAGATCGTAAATTAGATATTACAAGTCACAATGAAGATTACACGGTCGTAGAGCAGTATGAGCGCAGTGAGGGGCGTCACTCTACGGGCGGTATGGACGAGTTATACAAGTAA

>sfGFP
ATGCGTAAAGGCGAAGAGCTGTTCACTGGTGTCGTCCCTATTCTGGTGGAACTGGATGGTGATGTCAACGGTCATAAGTTTTCCGTGCGTGGCGAGGGTGAAGGTGACGCAACTAATGGTAAACTGACGCTGAAGTTCATCTGTACTACTGGTAAACTGCCGGTACCTTGGCCGACTCTGGTAACGACGCTGACTTATGGTGTTCAGTGCTTTGCTCGTTATCCGGACCATATGAAGCAGCATGACTTCTTCAAGTCCGCCATGCCGGAAGGCTATGTGCAGGAACGCACGATTTCCTTTAAGGATGACGGCACGTACAAAACGCGTGCGGAAGTGAAATTTGAAGGCGATACCCTGGTAAACCGCATTGAGCTGAAAGGCATTGACTTTAAAGAAGATGGCAATATCCTGGGCCATAAGCTGGAATACAATTTTAACAGCCACAATGTTTACATCACCGCCGATAAACAAAAAAATGGCATTAAAGCGAATTTTAAAATTCGCCACAACGTGGAGGATGGCAGCGTGCAGCTGGCTGATCACTACCAGCAAAACACTCCAATCGGTGATGGTCCTGTTCTGCTGCCAGACAATCACTATCTGAGCACGCAAAGCGTTCTGTCTAAAGATCCGAACGAGAAACGCGATCATATGGTTCTGCTGGAGTTCGTAACCGCAGCGGGCATCACGCATGGTATGGATGAACTGTACAAATGA

>ECK120029600
TTCAGCCAAAAAACTTAAGACCGCCGGTCTTGTCCACTACCTTGCAGTAATGCGGTGGACAGGATCGGCGGTTTTCTTTTCTCTTCTCAA

>L3S3P21**
GGCCAATTATTGAAGGCCTCCCTAACGGGGGGCCTTTTTTTGTTTCTGGTCTGCCCTC
